# Transcranial Direct current stimulation does not modulate performance on a tongue twister task

**DOI:** 10.1101/852483

**Authors:** Charlotte E. E. Wiltshire, Kate E. Watkins

**Affiliations:** Wellcome Centre for Integrative Neuroimaging, Department of Experimental Psychology, Radcliffe Observatory Quarter, University of Oxford, OX2 6GG, UK

**Keywords:** Non-invasive brain stimulation, Speech production, Articulatory motor cortex, Electrical stimulation, Speech motor learning

## Abstract

**Background:** TDCS modulates cortical excitability in a polarity-specific way. When used in combination with a behavioural task, it can also alter performance. Previously, tDCS modulated the performance of older adults on a complex speech motor learning task, which involved repetition of tongue twisters [1].

**Objective:** We aimed to replicate this finding in healthy young participants and to extend it by measuring tDCS-induced changes in motor excitability using transcranial magnetic stimulation and motor-evoked potentials elicited in the lips.

**Method:** In a double-blind randomized sham-controlled study, three groups of 20 participants received: 1) anodal tDCS to the left IFG/LipM1 and cathodal tDCS to the right hemisphere homologue; or 2) cathodal tDCS over the left and anodal over the right; or 3) sham stimulation. Participants heard and repeated tongue twisters and matched simple sentences before, during and 10 minutes after the stimulation. Motor excitability was measured before and immediately after the tDCS.

**Results:** The improvement in performance of tongue twister repetition from baseline to after stimulation was significantly greater than for the simple sentences but did not differ among the three groups. Motor excitability significantly decreased to a small but similar extent across the three groups.

**Conclusions:** TDCS did not modulate performance on a complex articulation task in healthy young adults. TDCS applied concurrently with task learning also failed to modulate motor excitability in expected ways. TDCS may be most effective in brains where brain function is sub-optimal due to age-related declines or pathology.

## Introduction

Transcranial direct current stimulation (tDCS) is a non-invasive brain stimulation technique that can modulate cortical excitability. tDCS exerts its effect by passing a weak electric current between two electrodes placed on the scalp. This current induces polarity-specific modulation such that the anode up-regulates, and the cathode down-regulates local cortical excitability [2,3]. The electrophysiological effects of tDCS can be demonstrated in the motor cortex by measuring changes in excitability via changes in the size of motor evoked potentials (MEPs) recorded from muscles of interest in response to single pulse transcranial magnetic stimulation (TMS) over the cortical representation [4]. Specifically, when targeting the representation of the hand in primary motor cortex (M1), the size of the MEP elicited by TMS in the contralateral hand muscles was increased by anodal tDCS (a-tDCS), and decreased by cathodal tDCS (c-tDCS) relative to sham stimulation [5,6]

When used in combination with a behavioural task, a single session of tDCS can modulate performance. For speech production specifically, there is some evidence to suggest that tDCS can modulate performance in neurologically intact speakers [7–9]. Previously, tDCS modulated performance on a task involving the repetition of tongue twisters [1]. Such sentences have complex articulation and their production often results in speech errors that are commonly seen in populations with speech pathologies [10]. Three groups of healthy older adults received a-tDCS or c-tDCS (both 2mA; 20 mins) or sham stimulation with the active electrode over the left IFG and the inert electrode over the contralateral frontopolar cortex. Participants’ response times and accuracy in repeating tongue twisters were successfully modulated during stimulation: a-tDCS significantly increased accuracy and reduced response times relative to baseline measures, whereas c-tDCS significantly reduced accuracy and increased response times from baseline and sham had no effect. A recent study failed to replicate this behavioural effect [11].

Here, we aimed to replicate the previously reported effects of tDCS on tongue twister repetition in healthy young adults and extended the design to include measures of changes in motor excitability before and after tDCS. In the current study, tDCS electrodes were placed over the ventral portion of motor cortex in each hemisphere encompassing the lip representation in M1, premotor and prefrontal cortex. Single pulses of transcranial magnetic stimulation were applied over the representation of the lips in the left M1 to elicit MEPs in the contralateral orbicularis oris muscle. This allowed us to gain more insight into the cortical effects induced by tDCS alongside behavioural outcomes in the same individuals, thus providing more sensitive information about the individual variability of cortical responses to tDCS [12,13]. The degree to which cortical excitability changes (as indexed by changes in MEP size) could predict changes in behaviour is unknown. Such a relationship could be an important consideration for the use of tDCS as a therapeutic tool. For example, participants could be tested for their suitability for tDCS treatments based on cortical measures of tDCS-induced modulation.

Our study is a conceptual replication in which we aimed to replicate the findings of a previous study [1] in a different population. It is not an exact replication as we made several important changes to the protocol (discussed below).

The study design and analysis plan were pre-registered on the Open Science Framework (https://osf.io/p84ys/).

## Methodology

### Sample size

To determine group sample sizes, we estimated an effect size based on graphs presented in [1] for comparing pairs of groups on the change in performance from baseline. We determined that 20 participants per group (n=60) were required based on a moderate effect size of Cohen’s d = 0.8 [1] with 80% power at the standard .05 alpha error probability (for a directional t-test). This sample size was double that of the previous study [1]; n=10 per group).

### Participants

Seventy-one healthy participants were recruited. They were all right-handed, native English speakers with normal or corrected-to-normal vision and normal hearing. We were unable to reliably elicit MEPs at a comfortable stimulation threshold in 11 participants. The remaining 60 participants were aged between 18 and 42 years (mean = 22.3, SD = 4.85); there were 30 men and 30 women.

The University of Oxford Central University Research Ethics Committee approved the study. Participants gave informed written consent to participate in the study, in accordance with the Declaration of Helsinki, and with the procedure approved by the committee.

### Design

The study was a double-blind randomized controlled study. Block randomization (block size 6) was used to assign 60 participants to one of four stimulation configurations with an allocation ratio of 2:1 (1mA tDCS with either anode on left or cathode on left, 20 participants each; sham with either anode on left or cathode on left, 10 participants each). Men and women were randomized separately to ensure equal genders in all groups. The researchers were blinded to the allocation of group by using the ‘study mode’ of the stimulator (NeuroConn GMbH, Ilmenau, Germany). A member of the research group who was not involved in the study assigned a 5-digit code to each participant. The link between the code and the stimulation group was not revealed until all 60 complete data sets were collected.

**Figure 1.**
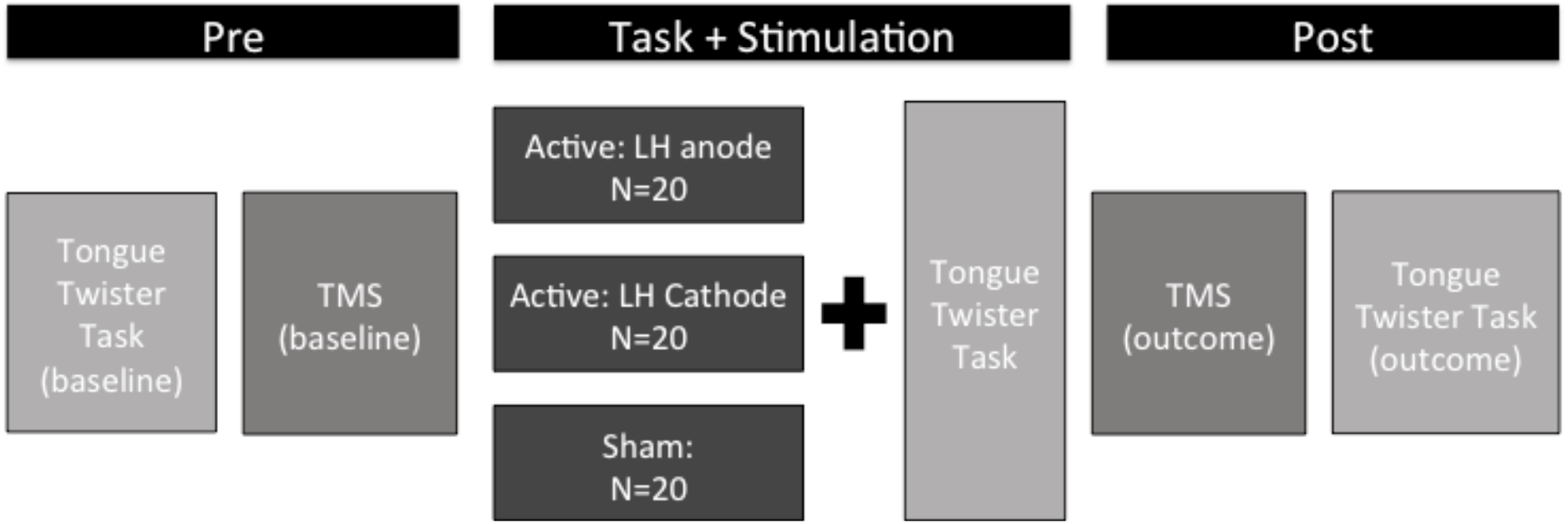
Study Design. The stimulation site and motor threshold were determined at baseline and 20 MEPs obtained. The tongue-twister task involved repetition of 36 tongue twisters and 36 simple sentences. The stimulation (anodal, cathodal and sham) was applied concurrently with the task. Twenty MEPs were obtained at the end of the stimulation period and the task was repeated again without stimulation.

### Procedure

#### Tongue Twister Task

36 novel tongue twisters (7 or 8 syllables) were taken from a previous published set [14]. For each tongue twister, a corresponding simple sentence was created that did not contain difficult articulation. To achieve this, one word was retained from the tongue twister and all other words were replaced by a word with the same number of syllables and syntactic class but without difficult articulation. For example, “Chad bravely wore Anne’s little shoes” was created to match the tongue twister “Brad bravely broke Brooke’s brittle blades”. A female, native-English speaker was recorded speaking the tongue twisters and control sentences in a soundproof booth. Recordings were presented via TMS-compatible insert headphones (Etymotic, Elk Grove Village, IL, USA).

For each trial, participants were instructed to repeat the sentence immediately after the cessation of the audio recording within a four and a half second response window. The task comprised three blocks of 24 sentences (12 tongue twisters and 12 simple sentences, presented in a randomised order). After each block, participants took a 30-s break. The duration of the task was 13 minutes. Participants practised the task on three simple sentences prior to the first run.

Response time was measured in the same way as previously [1], namely from the end of the stimulus presentation to the end of the participant’s spoken response. This measure encompasses both reaction time and total duration of the repeated sentence and included any hesitations, self-corrections or other types of dysfluency.

#### tDCS Stimulation

1mA of stimulation was delivered using a neuroConn GBH stimulator (NeuroConn GMbH, Ilmenau, Germany) via two 5 × 7cm saline-soaked electrodes. Two groups received active stimulation during which the intensity of current was ramped up slowly for 15 seconds before being held constant for 13 minutes and ramped down for 15 seconds. During the sham stimulation, the intensity of the current was ramped up slowly for 15 seconds before being held constant for 30 seconds and ramped down for 15 seconds. These sham stimulation parameters delivered current at an ineffective dosage [15].

The anodal group received bi-hemispheric, active stimulation with the anode placed on the left hemisphere IFG/M1 and the cathode placed over the homologous area in the right hemisphere. The cathodal group received bi-hemispheric, active stimulation with the reverse electrode configuration: the cathode over left hemisphere IFG/M1 and the anode over the homologous area in the right hemisphere. To ensure blinding of the researcher, placement of the electrodes was counterbalanced for the sham group such that half were placed in the anodal condition described above and half were placed in the cathodal configuration. A simulation of current density and flow based on the equipment and parameters used in this set up is illustrated in figure 2.

**Figure 2.**
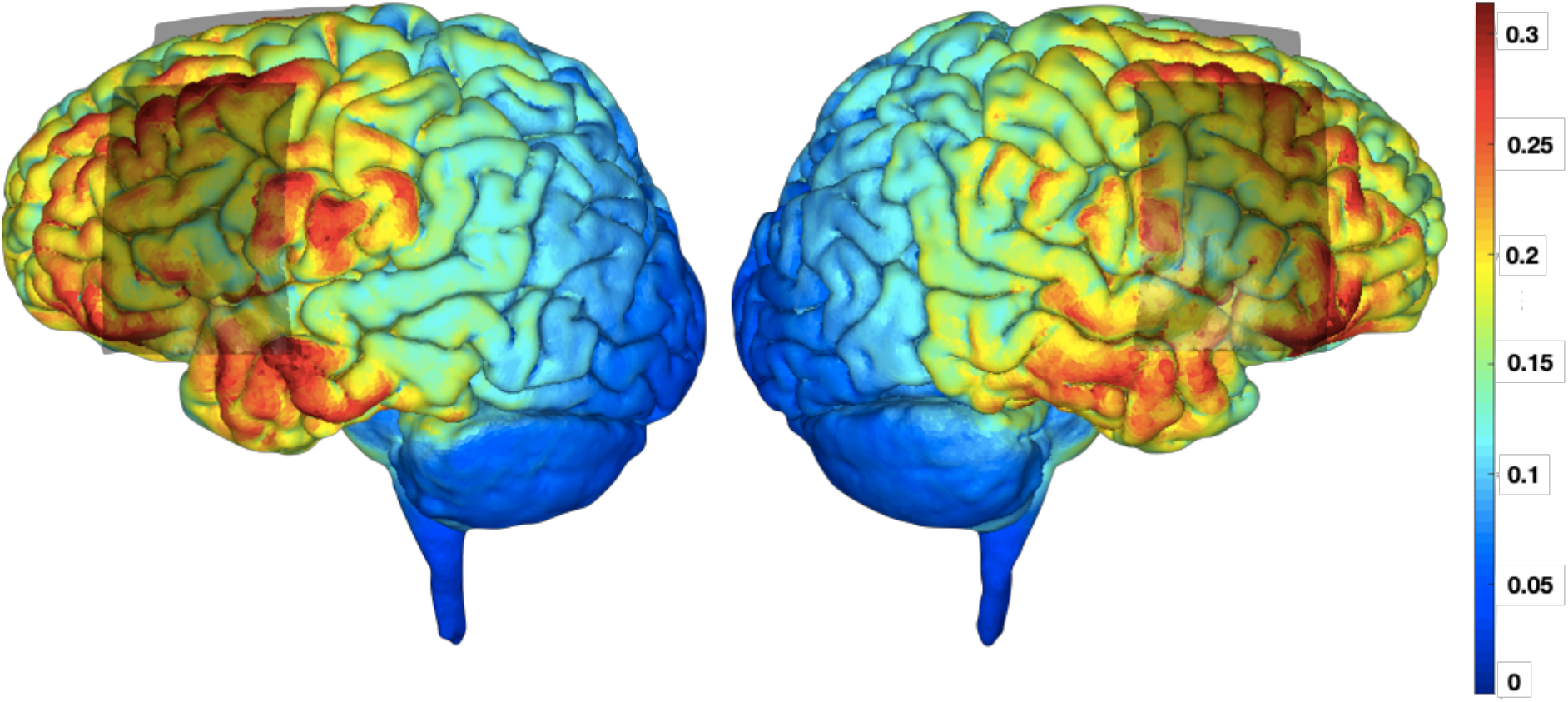
Simulated current flow. For this example, the anode is placed on the left hemisphere and the cathode on the right hemisphere. Red indicates high current density; blue indicates low current density. Simulation created using simnibs.com [16].

#### TMS and Electrophysiological recording

Single-pulse TMS was delivered using a DuoMag 200 stimulator through a 70-mm figure-eight coil. The coil was placed tangential to the skull, to induce a horizontal current flow from posterior to anterior under the junction of the two wings of the figure-eight coil. Surface electrodes (22 × 30 mm ABRO neonatal electrocardiogram electrodes) were attached to the right corner of the lower and upper lip (orbicularis oris muscle) in order to record the electrical activity of the underlying muscle. The ground electrode was attached to the forehead.

The active motor threshold was identified as the stimulation intensity needed to achieve an average MEP size that was at least 1mV peak-to-peak for 10 consecutive pulses whilst the participant maintained lip muscle contraction at 20% of their maximum. Subsequently, a train of 20 single pulses of stimulation were delivered with minimum 5-s intervals at this threshold to the left lip motor cortex to elicit the MEPs for measurement. Participants maintained the contraction of the lip muscle throughout the measurement.

For the MEP measures at post-stimulation, 20 MEPs were elicited using the same threshold (% of stimulator output) and position. BrainSight neuronavigation equipment (Rogue Research Inc, Montreal, Quebec, Canada) was used to ensure the precise area (accurate to 2 mm) is stimulated in an identical way (position, orientation and tilt of the coil) before and after the tDCS stimulation.

The peak-to-peak amplitude of the MEPs was calculated automatically within a window of 10-40 ms after the TMS pulse. The power of the rectified EMG signal for 200 ms before the TMS pulse was used to estimate the power of the contraction for each trial. As the amount of contraction is linearly related to the size of the MEP, we used analysis of covariance for each participant to adjust the MEP size for the amount of contraction (see [17]). This adjusted MEP size was used in the analyses below.

## Results

The results for each group are summarised in Table 1.

**Table 1.** Summary of participant demographics and results.

| Stimulation Group | Anodal |  |  | Cathodal |  |  | Sham |  |  |
| --- | --- | --- | --- | --- | --- | --- | --- | --- | --- |
| **Age**<br>*Mean (range; SD)* | 22.45 (19-33; 3.63) |  |  | 23.25 (19-42; 7.14) |  |  | 21.25 (18-29; 2.57) |  |  |
| **TMS output (% max output)**<br>*Mean (SEM)* | 56.6 (1.61) |  |  | 57.9 (2.22) |  |  | 62.2 (2.04) |  |  |
| Time point (relative to tDCS) | Pre | During | Post | Pre | During | Post | Pre | During | Post |
| **Tongue Twisters**<br>**Response time (s)**<br>*Mean (SEM)* | 3.25<br>(0.07) | 3.14<br>(0.06) | 3.08<br>(0.08) | 3.22<br>(0.08) | 3.01<br>(0.08) | 2.92<br>(0.09) | 3.13<br>(0.07) | 3.00<br>(0.08) | 2.93<br>(0.08) |
| **Simple Sentences**<br>**Response time (s)**<br>*Mean (SEM)* | 2.89<br>(0.07) | 2.77<br>(0.07) | 2.79<br>(0.09) | 2.80<br>(0.10) | 2.63<br>(0.09) | 2.56<br>(0.10) | 2.75<br>(0.07) | 2.61<br>(0.08) | 2.62<br>(0.08) |
| **Power of lip contraction (mV)**<br>*Mean (SEM)* | 0.117<br>(0.008) | NA | 0.116<br>(0.008) | 0.142<br>(0.112) | NA | 0.141<br>(0.011) | 0.122<br>(0.010) | NA | 0.120<br>(0.009) |
| **MEP size (mA)**<br>*Mean (SEM)* | 1.15<br>(0.05) | NA | 1.05<br>(0.05) | 1.16<br>(0.06) | NA | 1.13<br>(0.07) | 1.19<br>(0.07) | NA | 1.14<br>(0.08) |

SD = standard deviation, tDCS = transcranial direct current stimulation, SEM = standard error of the mean, mV = millivolts, mA = milliamps, NA = Not applicable.

Data will be made available on OSF. The following analyses were unchanged from the pre-registration analysis plan.

### Control Analyses

Firstly, the baseline data were analysed to confirm that the tongue twisters were repeated with longer durations compared with simple sentences (positive control) and also that there were no existing group differences at baseline. These data are plotted in Figure 3.

**Figure 3.**
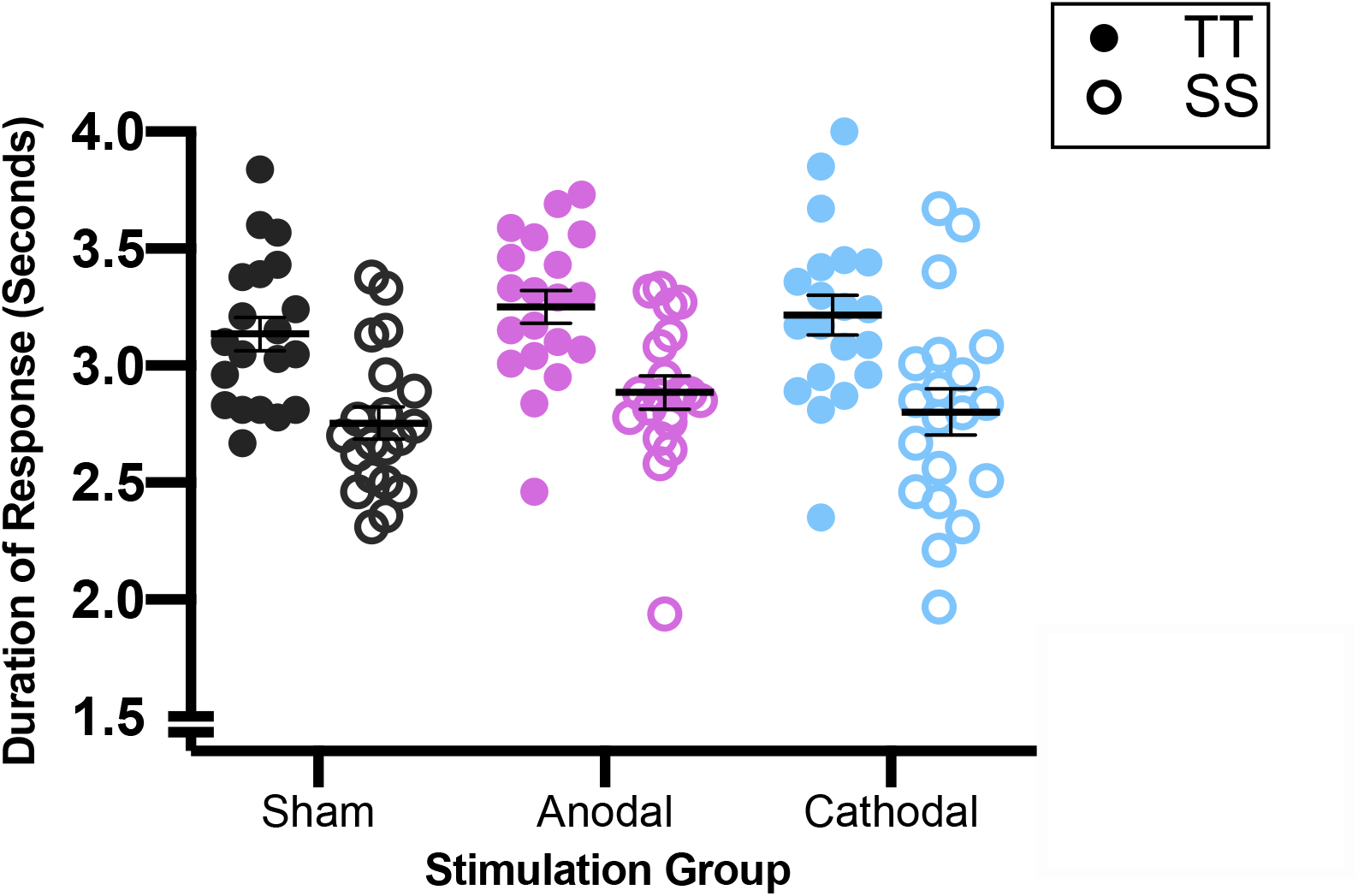
Baseline Task Performance. Duration of responses by sentence type (TT and SS) and stimulation group. Each point represents the mean of an individual participant. The horizontal black line is the group mean, error bars represent one SEM.

For the measure of response duration obtained pre-stimulation in the three groups, a 2 × 3 repeated measures analysis of variance (RM ANOVA), with sentence type as a within-subject factor (TT vs SS) and stimulation group as between-subject factor (anodal vs. cathodal vs. sham). There was a significant main effect of sentence type (F(1,57) = 509.72, *p* < .001, d = 5.4) due to longer response times for TT compared with SS in all three groups. There was no main effect of group (F(2,57) < 1) and no interaction between sentence type and group (F(2,57) < 1).

### Q1. Behavioural: Does anodal tDCS enhance learning to repeat tongue twisters in healthy young adults?

We used a 2 × 3 RM ANOVA with sentence type as a within-subject factor (TT, SS) and stimulation group as a between-subject factor (anodal, cathodal and sham) to test our hypotheses: H1A) people receiving anodal stimulation over the left hemisphere will show significantly greater improvements in sentence durations when repeating tongue twisters compared with people receiving cathodal or sham stimulation; H1B) people receiving cathodal stimulation over the left hemisphere will show significantly lower improvements in sentence durations when repeating tongue twisters compared with the people receiving sham stimulation; and H1C) the effect of anodal stimulation on sentence durations will be greater for repetition of tongue twisters compared with repetition of simple sentences. To assess learning, we analysed the dependent measure of change in duration over time (post-minus pre-stimulation). A significant main effect of sentence type showed the magnitude of reduction in response time was significantly greater for TT than for SS (F(1,57)=9.67, *p*=.003, d = 0.82). There was no main effect of group (F(2,57)=2.20, *p*=.120) or interaction between group and sentence type (F(2,57) < 1).

All groups showed a significant reduction in the duration of responses for repetition of sentences (i.e. one-sample t-test against no change: tongue twisters t(59) = −8.16, *p* <.001, d = 1.05; simple sentences t(59) = −4.80, *p* < .001, d = 0.62). This effect was significantly greater for the tongue twisters compared with the simple sentences, as shown in Figure 4. However, the lack of main effect or an interaction involving stimulation group indicates that task performance and this effect were not modulated by the anodal (or the cathodal) stimulation as predicted. Therefore, no further planned comparisons were carried out.

**Figure 4.**
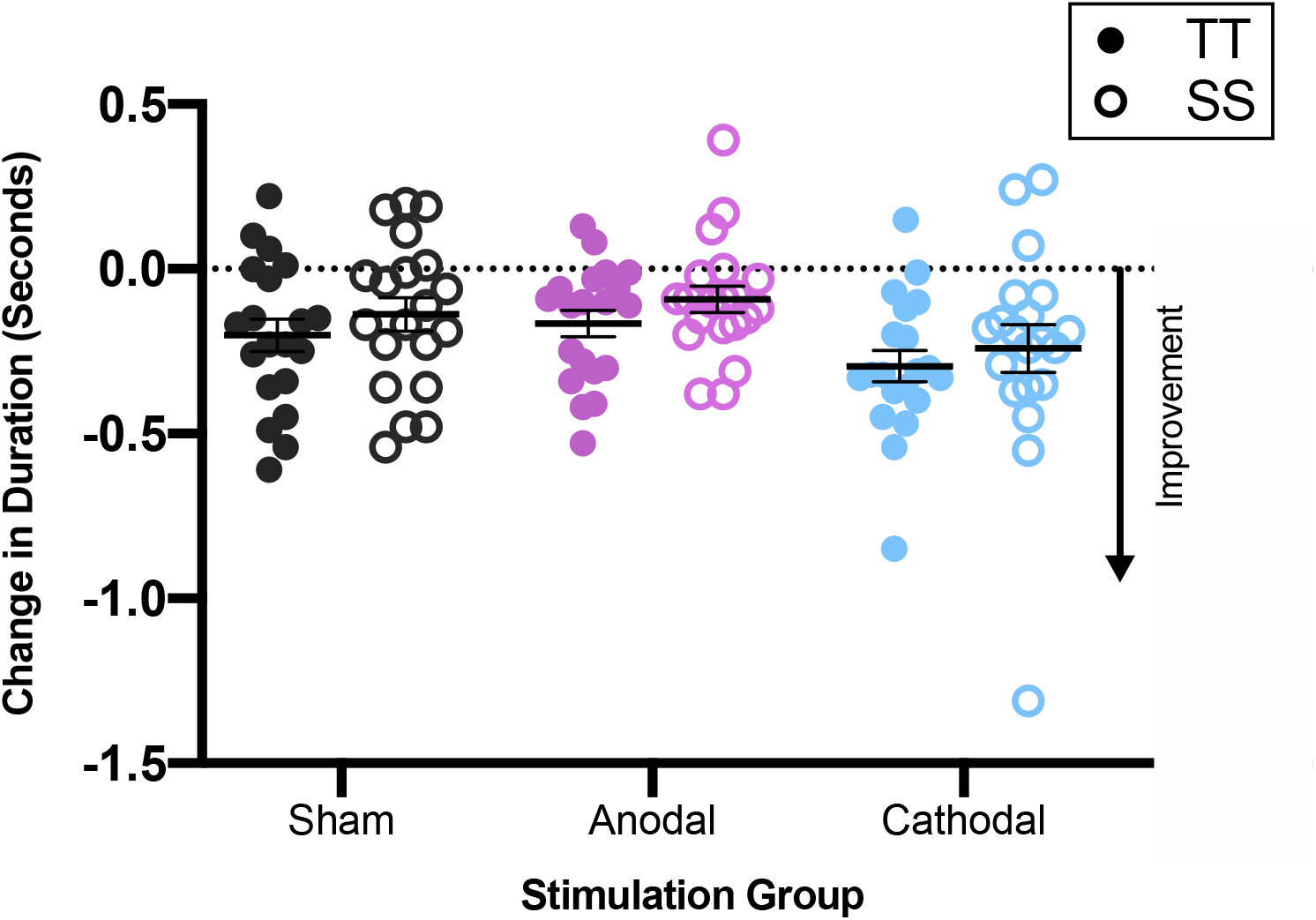
No Effect of tDCS on Learning to Repeat Tongue Twisters. Change in duration of responses (post- minus pre- stimulation) for tongue twisters (TT) and simple sentences (SS) by stimulation group. Each point represents the mean of an individual participant. Horizontal black line shows group mean. Error bars represent SEM.

### Q2. Electrophysiological: Does tDCS change excitability in the motor system underlying speech production?

To test our Hypothesis 2A (anodal tDCS over the left hemisphere will increase excitability in the speech motor system measured contralaterally compared with cathodal and sham stimulation), two t-tests compared change in MEP size (post- minus pre- stimulation) between the anodal and cathodal groups and separately between the anodal and sham groups. Directional t-tests were used as we expected an increase in MEP amplitude in the anodal group compared with the other two groups, which we expected to remain either unchanged (sham) or to decrease in amplitude (cathodal group). The change in MEP size for the anodal group was not significantly bigger than the changes for either of the other two groups (anodal vs sham: t(38) = 0.80, *p* = .469; anodal vs cathodal: t(38) = −0.31, *p* = .380).

To test our Hypothesis 2B (cathodal tDCS over the left hemisphere will decrease excitability in the speech motor system measured contra-laterally compared with sham stimulation) a single t-test was used to compared change in MEP size between cathodal and sham groups. A directional test was used as we expected a decrease in MEP amplitude in the cathodal group compared with the sham group, which we expected would not change. The change in MEP size for the cathodal group was not significantly different from that in the sham group (t(38) = 0.29, *p* = .387).

**Figure 5.**
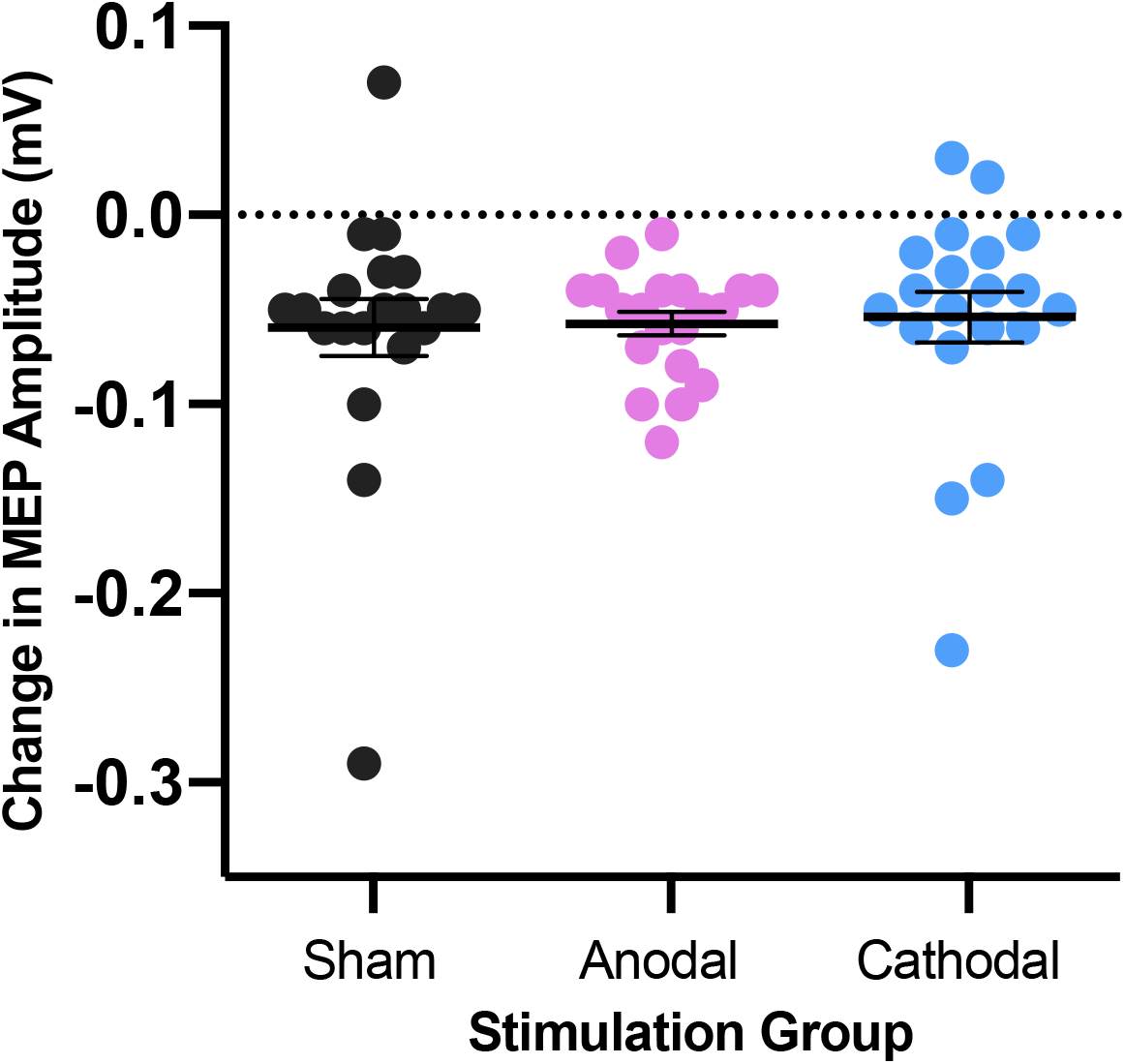
No Effect of tDCS on Excitability in the Motor System Underlying Speech Production. Change in MEP size (post- minus pre- stimulation) by stimulation group. Each point represents the mean of an individual participant. Horizontal black line shows group mean. Error bars represent SEM. Dashed line at y=0 represents no change in MEPs.

### Q3. Does the change in motor excitability predict learning on the behavioural task?

To test our Hypothesis 3A that change in motor excitability will correlate positively with the size of the improvement on the tongue twister task we correlated the change in MEP size with the change in duration of repetition of tongue twisters. The change in MEP size did not correlate with change in duration of repetition of tongue twisters for any of the groups (Anodal: r = −.33, *p* = .161; Cathodal: r = −.07, *p* = .757; Sham: r = −.14, *p* = .517). We also planned to compare the slopes of the regression lines in the three groups separately. However, because none of the correlations were significant, these comparisons were not carried out.

### Exploratory analysis

The following analyses were not planned.

### Did Task Performance Change During Stimulation?

We also tested whether tDCS affected task performance during stimulation as it did in the previous study [1]. The means and SEM for each group are shown in Table 1.

The change in duration of response from pre- to during-stimulation was significantly different from zero (no change) for all groups (all *p* < .005), however neither anodal nor cathodal were different from sham (anodal vs sham: t(38) = −.64, *p* = .529; cathodal vs sham: t(38) = 1.09, *p* = .284). All groups improved their performance but tDCS had no effect on this improvement.

### Did the Amount of Muscle Contraction During MEP Measurements Differ Pre- and Post-tDCS?

Differences in the power of the lip muscle contraction during TMS affects MEP size. We therefore tested whether this had changed from pre- to post-tDCS in any of our groups (see Table 1). There were no differences between the power of contraction at pre- and post-tDCS for any of the stimulation groups (sham: t(19) = 0.47, *p* = .645; anodal: t(19)= 0.859, *p* = .401; cathodal: t(19) = 0.29, *p* = .777).

MEP amplitudes significantly decreased from pre- to post-stimulation for all of the groups (see Fig. 6) (one-sample t-test against no change, i.e. zero: t(59) = −8.24, *p* < .001, d = 1.06).

### Were participants blind to whether they were receiving real or sham stimulation?

In order to test whether participants could guess if they were receiving real or sham stimulation, a 2 × 2 Chi-square test was performed. Responses from seven participants were not recorded (sham = 3, real = 4). Of the 17 people who received sham, 8 guessed it was sham and 9 that it was real stimulation; of the 36 people who received real stimulation, 11 guessed it was same and 25 that it was real. These proportions are not different to chance (χ^2^(1, N = 53) = 1.37, *p* = .242) indicating that participants were successfully blinded to the type of stimulation they were receiving.

## Discussion

We tested whether tDCS could modulate performance on a tongue-twister task in healthy young adults. Sixty participants received either sham (n=20), or bi-hemispheric tDCS with the anode on the left (n=20) or right (n=20). Their ability to repeat sentences that were either complex (tongue twisters) or simple was tested before and after tDCS concurrent with the task. TDCS did not modulate performance on the task. Participants showed an improvement in performance and this was greater for the tongue twisters than for the simple sentences but these changes did not differ among the three groups. These results align with a recent study that also failed to replicate the original behavioural finding [11]. The effect of tDCS on behaviour in neurotypical populations has been difficult to replicate [13], [18]. This could in part reflect individual differences in the expected response to brain stimulation. Therefore, we also used TMS to elicit MEPs as a measure of motor excitability before and after the tDCS; we predicted that motor excitability would be modulated by tDCS in a polarity-specific way and that individual differences in the behavioural effects of tDCS might be explained by differences in the change in motor excitability. Our results showed no modulatory effect of tDCS on motor excitability. Participants showed a small reduction in excitability from pre- to post-stimulation but this did not differ among the three groups.

These results contrast with those previously reported [1]. There are a number of differences between study protocols that might explain the failure to replicate. Firstly, we changed the electrode montage from uni-hemispheric to bi-hemispheric and placed them slightly more posteriorly in order to ensure stimulation of the lip representation of M1. It is unlikely that the slight difference in position of large electrodes compared with the original study reduced the effectiveness of tDCS coupled with the task as indicated by our modelling of the current flow for our study (see Fig. 2). In addition, bi-hemispheric montages are at least as effective as uni-hemispheric ones [19–21] or can even improve the effects on task performance [22–24]. Secondly, we reduced the current amplitude from 2mA to 1mA and the duration of stimulation from 20 minutes to 13 minutes. In our experience and that of other reports, blinding of the participant is not achieved at 2mA due to the increased somatosensory experiences, e.g. tingling and itching under the electrodes [25]. Our debriefing indicated that participants were blind to stimulation type at 1mA. Previous reports [26] and our own experience [27] suggests that increasing stimulation intensity beyond 1mA does not increase the effectiveness of anodal stimulation over the ventral motor cortex. In addition, reducing the duration of stimulation is unlikely to explain our null results as numerous reports of tDCS applied to the motor cortex in healthy humans show behavioural modulation with stimulation durations of between 10 and 16 minutes [4,6,7,28,29]. In sum, we believe the changes to the stimulation protocol described above are unlikely explanations for our failure to detect a modulatory effect on task performance in this study. We turn next to the changes we made to the behavioural protocol.

Our study used different stimuli for the task and introduced a control task (repetition of simple sentences). Necessarily, the language of the sentences was changed from Italian to English. If the speech motor effect is generalisable across languages, this would not explain the difference in results. Our tongue twisters were novel rather than well-known (as in [1]) and shorter than those used previously; the mean response time for tongue twisters at baseline in our study was 3.2 seconds, whereas it was ~4.5 seconds in the previous study [1]. Our study may have been less sensitive to changes in behaviour between stimulation groups because of these differences in stimuli.

The most important difference between the two studies was the age of the participants. Both studies included neurotypical participants but those in [1] were considerably older (mean = 57 years; SD = 11) than those in the current study (mean = 22.3 years, SD = 4.8). In our view, this age difference is the most plausible explanation for the different findings between the two studies. For example, in a previous study, TDCS with a concurrent visuomotor adaptation task significantly improved performance of healthy older adults to the level of that seen in younger adults without stimulation [30] suggesting that age-related declines in task performance can be reversed using tDCS. Taken together, the reduction in the sentence length for our study and our focus on younger healthy adults may have reduced our sensitivity to the modulatory effects of tDCS [31].

In the current study, we added electrophysiological measurements of motor excitability to assess tDCS changes using TMS-induced MEPs. Our aim was to explain individual differences in the anticipated modulatory effects of tDCS on task performance by variability in the modulatory effects of tDCS on motor excitability. In some respects, we succeeded, in that the failure to find an effect of tDCS on task performance was mirrored by a lack of effect of tDCS on motor excitability. Nevertheless, this result was unexpected given previous established results that tDCS modulates MEP size in a polarity-specific way [3]. It is important to note, however, that these modulatory effects were found in studies that did not involve a concurrent task [3,4]. A previous report found that anodal tDCS (1mA, 20 minutes) applied without a concurrent task increased MEP size as expected but when the stimulation was applied in combination with a digit-sequence task, MEP size was not modulated even though task performance measurably improved [32]. One explanation of these results is that during task performance the brain alters its excitability to counteract the effects of tDCS. Such homeostatic regulation would therefore abolish the measurable effects of tDCS on motor excitability. This suggestion could be important in explaining the variable results within the tDCS literature and aid optimisation of tDCS protocols.

It is possible that tDCS is most effective when the area of cortex being stimulated functions atypically. For example, left ventral premotor cortex is known to be underactive during speaking in people who stutter compared with controls [33] and anodal tDCS over this area improved fluency in people who stutter compared with sham stimulation [34]. Similarly, tDCS led to modulated performance on a digit sequence task in the non-dominant, but not the dominant hand of neurotypical adults (Boggio et al., 2006). In our opinion, the negative results for both task and motor excitability in the current study are best explained by the fact that our healthy young adults function optimally, which renders modulation by tDCS ineffective. Note, that this is not simply due to a behavioural ceiling effect as there was room for improvement on task performance both in terms of latency and accuracy, which would have affected response time. Furthermore, the cathodal stimulation was expected to lower performance and was also ineffective.

In summary, our study failed to demonstrate the previously reported polarity-specific modulatory effects of tDCS on speech motor control in a typical population. Our study had a sample size double that of the previous study and was sufficiently powered to detect a similarly sized effect. The factor of participant age and how this interacts with brain function is the most likely explanation for this failure to detect an effect should one exist. The alternative explanation is that the effect cannot be replicated but the changes we made to the protocol and the population difference in age precludes such a firm conclusion. The lack of modulation by tDCS on motor excitability is consistent with the lack of effect on behaviour but we believe this is better explained by homeostatic regulation of cortical excitability that may occur during task performed concurrently with tDCS in the typically functioning brain.

## References

[1] Fiori V, Cipollari S, Caltagirone C, Marangolo P. “If two witches would watch two watches, which witch would watch which watch?” tDCS over the left frontal region modulates tongue twister repetition in healthy subjects. Neuroscience 2014;256: 195–200. doi:10.1016/j.neuroscience.2013.10.048.

[2] Bikson M, Rahman A. Origins of specificity during tDCS: Anatomical, activity-selective, and input-bias mechanisms. Front Hum Neurosci 2013;7:688. doi:10.3389/fnhum.2013.00688.

[3] Nitsche MA, Paulus W. Excitability changes induced in the human motor cortex by weak transcranial direct current stimulation. J Physiol 2000;527:633–9. doi:10.1111/j.1469-7793.2000.t01-1-00633.x.

[4] Stagg CJ, Nitsche MA. Physiological Basis of Transcranial Direct Current Stimulation. Neurosci 2011;17:37–53. doi:10.1177/1073858410386614.

[5] Stagg CJ, O’Shea J, Kincses ZT, Woolrich M, Matthews PM, Johansen-Berg H. Modulation of movement-associated cortical activation by transcranial direct current stimulation. Eur J Neurosci 2009;30:1412–23. doi:10.1111/j.1460-9568.2009.06937.x.

[6] Nitsche MA, Schauenburg A, Lang N, Liebetanz D, Exner C, Paulus W, et al. Facilitation of implicit motor learning by weak transcranial direct current stimulation of the primary motor cortex in the human. J Cogn Neurosci 2003;15:619–26. doi:10.1162/089892903321662994.

[7] Lametti DR, Smith HJ, Freidin PF, Watkins KE. Cortico-cerebellar networks drive sensorimotor learning in speech. J Cogn Neurosci 2018;30:540–51. doi:10.1162/jocn_a_01216.

[8] Buchwald A, Calhoun H, Rimikis S, Lowe MS, Wellner R, Edwards DJ. Using tDCS to facilitate motor learning in speech production: The role of timing. Cortex 2019;111:274–85. doi:10.1016/j.cortex.2018.11.014.

[9] Deroche MLD, Nguyen DL, Gracco VL. Modulation of Speech Motor Learning with Transcranial Direct Current Stimulation of the Inferior Parietal Lobe. Front Integr Neurosci 2017;11. doi:10.3389/fnint.2017.00035.

[10] Wilshire CE. The “tongue twister” paradigm as a technique for studying phonological encoding. Lang Speech 1999;42:57–82. doi:10.1177/00238309990420010301.

[11] Wong MN, Chan Y, Ng ML, Zhu FF. Effects of transcranial direct current stimulation over the Broca’s area on tongue twister production. Int J Speech Lang Pathol 2019;21:182–8. doi:10.1080/17549507.2017.1417480.

[12] Chew T, Ho KA, Loo CK. Inter- and intra-individual variability in response to transcranial direct current stimulation (tDCS) at varying current intensities. Brain Stimul 2015;8:1130–7. doi:10.1016/j.brs.2015.07.031.

[13] Guerra A, López-Alonso V, Cheeran B, Suppa A. Variability in non-invasive brain stimulation studies: Reasons and results. Neurosci Lett 2018. doi:10.1016/j.neulet.2017.12.058.

[14] Fossett TRD, McNeil MR, Pratt SR, Tompkins CA, Shuster LI. The effect of speaking rate on serial-order sound-level errors in normal healthy controls and persons with aphasia. Aphasiology 2016;30:74–95. doi:10.1080/02687038.2015.1063581.

[15] Jog M V., Smith RX, Jann K, Dunn W, Lafon B, Truong D, et al. In-vivo imaging of magnetic fields induced by Transcranial Direct Current Stimulation (tDCS) in human brain using MRI. Sci Rep 2016;6. doi:10.1038/srep34385.

[16] Saturnino G, Antunes A, Stelzer J, Thielscher A. SimNIBS: A versatile toolbox for simulating fields generated by transcranial brain stimulation 2015.

[17] Watkins KE, Strafella AP, Paus T. Seeing and hearing speech excites the motor system involved in speech production. Neuropsychologia 2003;41:989–94. doi:10.1016/S0028-3932(02)00316-0.

[18] Tremblay S, Larochelle-Brunet F, Lafleur L-P, El Mouderrib S, Lepage J-F, Théoret H. Systematic assessment of duration and intensity of anodal transcranial direct current stimulation on primary motor cortex excitability. Eur J Neurosci 2016;44:2184–90. doi:10.1111/ejn.13321.

[19] Prichard G, Weiller C, Fritsch B, Reis J. Effects of different electrical brain stimulation protocols on subcomponents of motor skill learning. Brain Stimul 2014;7:532–40. doi:10.1016/j.brs.2014.04.005.

[20] Meinzer M, Lindenberg R, Sieg MM, Nachtigall L, Ulm L, Flöel A. Transcranial direct current stimulation of the primary motor cortex improves word-retrieval in older adults. Front Aging Neurosci 2014;6. doi:10.3389/fnagi.2014.00253.

[21] Fiori V, Nitsche M, Iasevoli L, Cucuzza G, Caltagirone C, Marangolo P. Differential effects of bihemispheric and unihemispheric transcranial direct current stimulation in young and elderly adults in verbal learning. Behav Brain Res 2017;321:170–5. doi:10.1016/j.bbr.2016.12.044.

[22] Vines BW, Cerruti C, Schlaug G. Dual-hemisphere tDCS facilitates greater improvements for healthy subjects’ non-dominant hand compared to uni-hemisphere stimulation. BMC Neurosci 2008;9. doi:10.1186/1471-2202-9-103.

[23] Drummond NM, Hayduk-Costa G, Leguerrier A, Carlsen AN. Effector-independent reduction in choice reaction time following bi-hemispheric transcranial direct current stimulation over motor cortex. PLoS One 2017;12. doi:10.1371/journal.pone.0172714.

[24] Waters S, Wiestler T, Diedrichsen J. Cooperation not competition: Bihemispheric tDCS and fMRI show role for ipsilateral hemisphere in motor learning. J Neurosci 2017;37:7500–12. doi:10.1523/JNEUROSCI.3414-16.2017.

[25] O’Connell NE, Cossar J, Marston L, Wand BM, Bunce D, Moseley GL, et al. Rethinking Clinical Trials of Transcranial Direct Current Stimulation: Participant and Assessor Blinding Is Inadequate at Intensities of 2mA. PLoS One 2012;7. doi:10.1371/journal.pone.0047514.

[26] Kidgell DJ, Daly RM, Young K, Lum J, Tooley G, Jaberzadeh S, et al. Different current intensities of anodal transcranial direct current stimulation do not differentially modulate motor cortex plasticity. Neural Plast 2013;2013. doi:10.1155/2013/603502.

[27] Chesters J. Enhancing Speech Fluency using Transcranial Direct Current Stimulation. University of Oxford, 2016.

[28] Monte-Silva K, Kuo MF, Hessenthaler S, Fresnoza S, Liebetanz D, Paulus W, et al. Induction of late LTP-like plasticity in the human motor cortex by repeated non-invasive brain stimulation. Brain Stimul 2013;6:424–32. doi:10.1016/j.brs.2012.04.011.

[29] Lang N, Nitsche MA, Sommer M, Tergau F, Paulus W. Chapter 28 Modulation of motor consolidation by external DC stimulation. Suppl Clin Neurophysiol 2003;56:277–81. doi:10.1016/S1567-424X(09)70231-4.

[30] Panouillères MTN, Joundi RA, Brittain JS, Jenkinson N. Reversing motor adaptation deficits in the ageing brain using non-invasive stimulation. J Physiol 2015;593:3645–55. doi:10.1113/JP270484.

[31] Fiori V, Nitsche MA, Iasevoli L, Cucuzza G, Caltagirone C, Marangolo P. Differential effects of bihemispheric and unihemispheric transcranial direct current stimulation in young and elderly adults in verbal learning. Behav Brain Res 2017;321:170–5. doi:10.1016/j.bbr.2016.12.044.

[32] Amadi U, Allman C, Johansen-Berg H, Stagg CJ. The homeostatic interaction between anodal transcranial direct current stimulation and motor learning in humans is related to GABAa activity. Brain Stimul 2015;8:898–905. doi:10.1016/j.brs.2015.04.010.

[33] Watkins KE, Smith SM, Davis S, Howell P. Structural and functional abnormalities of the motor system in developmental stuttering. Brain 2008;131:50–9. doi:10.1093/brain/awm241.

[34] Chesters J, Möttönen R, Watkins KE. Transcranial direct current stimulation over left inferior frontal cortex improves speech fluency in adults who stutter. Brain 2018;141:1161–71. doi:10.1093/brain/awy011.

